# Deep learning-based prediction of enzyme optimal pH and design of point mutations to improve acid resistance

**DOI:** 10.1101/2024.11.16.623957

**Authors:** Sizhe Qiu, Nan-Kai Wang, Yishun Lu, Jin-Song Gong, Jin-Song Shi, Aidong Yang

## Abstract

An accurate deep learning predictor of enzyme optimal pH is essential to quantitatively describe how pH influences the enzyme catalytic activity. CatOpt, developed in this study, outperformed existing predictors of enzyme optimal pH (RMSE=0.833 and R2=0.479), and could provide good interpretability with informative residue attention weights. The classification of acidic and alkaline enzymes and prediction of enzyme optimal pH shifts caused by point mutations showcased the capability of CatOpt as an effective computational tool for identifying enzyme pH preferences. Furthermore, a single point mutation designed with the guidance of CatOpt successfully enhanced the activity of *Pyrococcus horikoshii* diacetylchitobiose deacetylase at low pH (pH=4.5/5.5) by approximately 7%, suggesting that CatOpt is a promising *in-silico* enzyme design tool for pH-dependent enzyme activities.

**Graphical abstract:** 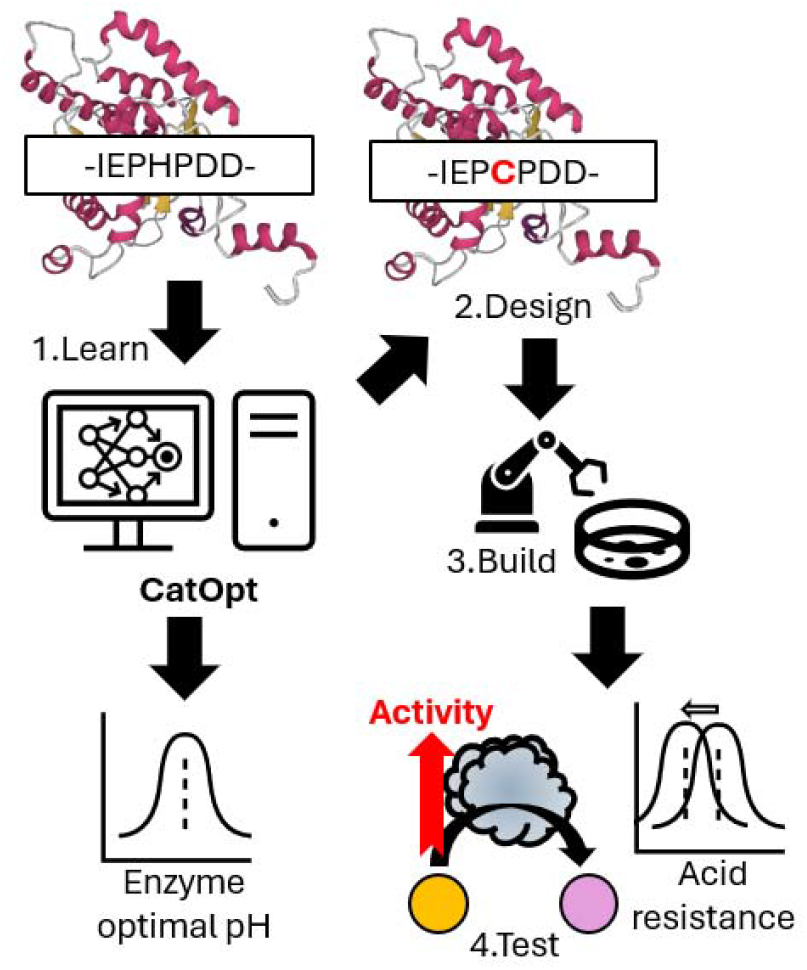

## 1. Introduction

In the era of synthetic biology, enzymes play a crucial role in industrial processes, such as food fermentation, waste transformation, and eco-friendly bio-manufacturing of chemical products [1]. In those industrial processes, pH is an important influencing factor of enzyme catalytic activity, as the increase or decrease of pH can affect enzyme protein conformations [2– 4]. Each enzyme has an optimal pH (*pH*_*opt*_) where its maximum catalytic rate is attained. Therefore, an accurate enzyme *pH*_*opt*_ predictor is highly desirable for enzyme mining and engineering, enabling the discovery of enzymes suited to specific environmental pH and supporting the efforts to enhance catalytic activity within targeted pH ranges.

To fill the knowledge gap of enzyme *pH*_*opt*_ in enzyme databases (e.g., BRENDA [5], uniprot [6]) caused by the high cost of enzyme assays [7], several machine learning models were developed to make predictions from protein sequences, but most of them could only predict *pH*_*opt*_ ranges (acidic or alkaline) [8–11]. MeTarEnz [12] used random forest regression to quantitatively predict *pH*_*opt*_ values, but the accuracy was low (MSE=1.648 and R2=0.195). Recently, the use of protein language models improved the prediction accuracy of *pH*_*opt*_. EpHod [13] and Seq2pHopt [14] achieved RMSE scores close to 1 using ESM-1 [15] and ESM-2 [16], respectively. Subsequently, OphPred [17] surpassed EpHod and Seq2pHopt using ESM-2 and XGBoost [18], but it lacked interpretability for protein residues. Besides, none of the existing models has been applied to enzyme engineering.

With the aim to build a predictor of enzyme *pH*_*opt*_ with good accuracy and interpretability, this study constructed a deep learning model, named CatOpt, with a pre-trained language model of proteins, multi-scale convolutional neural network (CNN), multi-head self-attention, and residual dense neural networks. CatOpt allows the interpretation of residue attention weights, which helps to decipher the key sequence information for enzyme *pH*_*opt*_. Case studies on classifying acidic and alkaline enzymes and predicting enzyme *pH*_*opt*_ changes by point mutations were carried out to examine CatOpt’s performance on *in-silico* enzyme selection. The predictor-guided engineering of *Pyrococcus horikoshii* diacetylchitobiose deacetylase to enhance acid resistance, suited to its acidic working environment [19], demonstrated that CatOpt can function as a useful computational tool for enzyme engineering.

## 2. Methods

### 2.1 Construction of the deep learning model

The training and test datasets of optimal pH (*pH*_*opt*_) were obtained from the Zenodo repository of EpHod [13]. The training set in this study was merged with the original training and validation sets of EpHod. The model architecture of CatOpt consisted of protein sequence embedding by ESM-2 [16], multi-scale convolutional neural network (CNN), multi-head self-attention network, and residual dense blocks (**Figure 1**). Using the same training and test datasets as EpHod avoided data leakage in model comparison (**section 3.1**).

**Figure 1.**
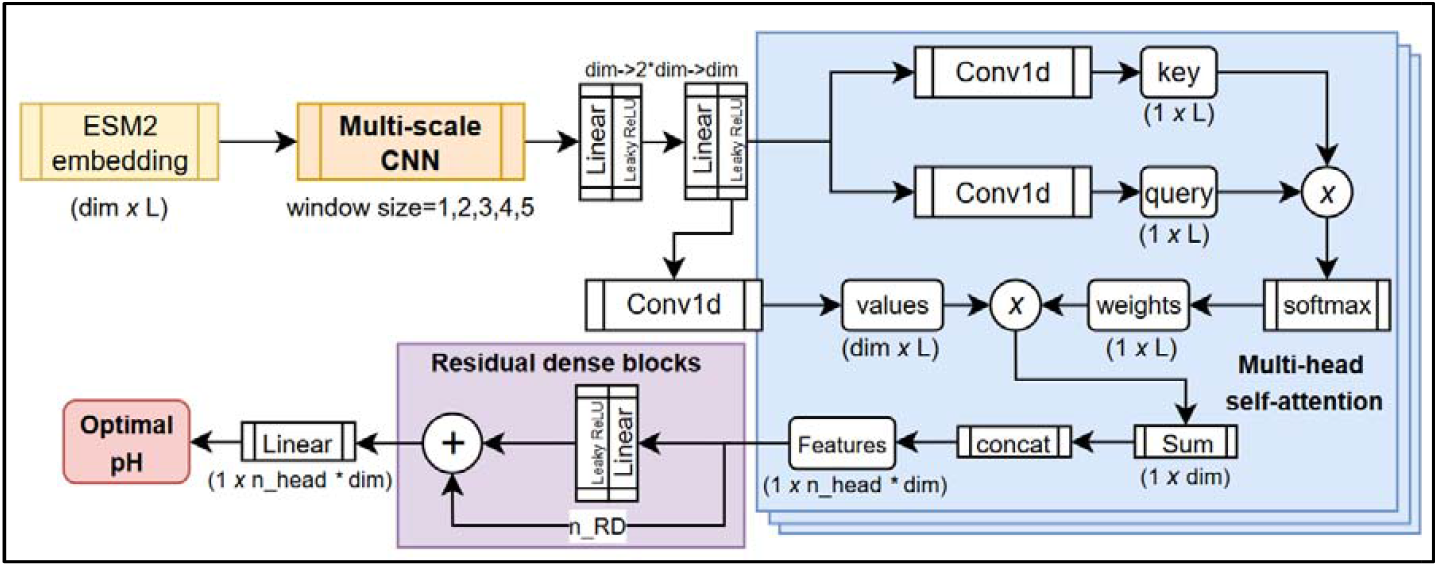
The model architecture of CatOpt. L: protein sequence length; dim: embedding dimension size; Conv1d: 1-D convolutional layer; ⊗: element-wise multiplication; n_head: the number of heads in multi-head attention; concat: concatenation; RD: residual dense block, a dense layer with residual connection.

First, the protein sequence embeddings (*r* ∈ *R*^*dim*×*L*^, *L*: *sequence length,dim* = 320) were computed using the esm2_t6_8M_UR50D model [16]. The embeddings (*r*) were first passed to the multi-scale CNN to extract features with sliding window sizes of 1, 2, 3, 4 and 5. The outputs from CNNs with different sliding window sizes were added in an element-wise way. Then, the outputs were passed to a 2-layer linear featuremap. The feature dimension was transformed from dim to 2*dim, and then back to dim.

In multi-head self-attention, the outputs of the 2-layer linear featuremap were transformed to values (*v*^*i*^ ∈ *R*^*dim*×*L*^, *L*: *attention head index*), keys (*k*^*i*^ ∈ *R*^1×*L*^) and queries (*q*^*i*^ ∈ *R*^1×*L*^) via 1-D CNNs. The self-attention weights (*w*^*i*^ ∈ *R*^1×*L*^) were computed with element-wise multiplication of keys and queries and a softmax function (Eq. 1). Next, the element-wise products of values and weights were computed and summed at the dimension of sequence length. The weighted features () from all attention heads were concatenated as the inputs () for residual dense blocks. Each residual dense block consisted of a linear layer, Leaky ReLU [20], and a residual addition operator, ⊕. In the end, a linear layer used the outputs from residual dense blocks to regress for enzyme values.

### 2.2 Deep learning model training

For the training process, batch training was used (batch size=32) for the efficiency and generalizability of the deep learning neural network. Adam optimization algorithm [21] was used to update neural network weights iteratively. The loss function was mean squared error (MSE). The initial learning rate was 0.0005, and the learning rate decayed by 50% for every 10 epochs to prevent overfitting. Before model training started, 10% of the training set was split out as the validation set, and target values were rescaled as 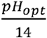. During the training process, the prediction accuracy of the model was evaluated with root mean squared error (RMSE), mean average error (MAE), and r-squared (R2) (**see SI, Eq. S1-3**). For details of software and hardware, please see the section S1.1 of the supplementary information. There were 2 hyperparameters in CatOpt, number of attention heads (n_head) and number of residual dense blocks (n_RD), and hyperparameter optimization was performed on n_head=4,5,6 and n_RD=3,4,5 (**see SI, Figure S2**).

### 2.3 Interpretation of residue attention weights

To investigate how enzyme *pH*_*opt*_ was predicted from the amino acid sequence, the average residue attention weights (*w*_*avg*_ ∈ *R*^1\**L*^) were computed by averaging the weights across all attention heads (Eq. 2). Then, the average residue attention weights were mapped to the protein sequence, together with annotated acidic/basic residues, active and binding sites obtained from the uniprot database [6]. The spatial distribution of residue attention weights and annotated protein sequence features could assist in revealing the key sequence information influencing the enzyme *pH*_*opt*_.

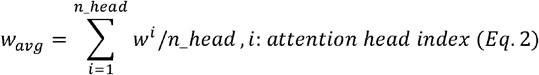

### 2.4 Predictor-guided design of single point mutations

CatOpt was used to design single point mutations to enhance the acid resistance of *Pyrococcus horikoshii* diacetylchitobiose deacetylase (PhDac), an enzyme catalyzing the production of glucosamine (GlcN) from N-acetylglucosamine (GlcNAc) [19]. Because the fermentation environment of diacetylchitobiose deacetylase is usually acidic, it is desirable to enhance its enzyme activity under low pH [22]. For 3*21 sites centered at three substrate binding sites (D46, R92, and H152, -10 site to +10 site), all possible amino acid substitutions were considered, and thus there were totally 1197 mutated sequences. For all mutants, *pH*_*opt*_ and turnover numbers were predicted by CatOpt and DLTKcat [23], respectively. DLTKcat was used to predict turnover numbers of mutants and filter out mutants with low catalytic efficiency. Designed point mutations were selected with an arbitrary threshold of predicted *pH*_*opt*_<7. In this study, site directed mutagenesis, protein expression and purification of PhDac mutants followed the same procedure as in Huang et al., 2021 [19].

### 2.5 Measurement of enzyme activity

The enzyme activity measurement method was adapted from Jiang et al., 2019 [24] with several modifications. In summary, the reaction solution comprised 1 mL of citrate buffer (50 mM, pH 4.5 to 5.5), 50 g/L of GlcNAc, and 100 µL crude enzyme (i.e., designed mutants). The reaction was performed in a metal bath at 40 °C and 900 rpm for 20 minutes. Subsequently, the reaction was halted by the addition of 50 µL of 0.5 M HCl. The mixture was then centrifuged at 12,000 rpm for 5 minutes. After centrifugation, 10 μL of the supernatant was combined with 100 μL of the OPA detection reagent (composed of 5 mg OPA, 10 μL of 1.0 M dithiothreitol, and 100 μL of alcohol in 10 mL of sodium carbonate buffer), and the absorbance was measured at 330 nm using the Infinite M200 PRO Spectrum spectrophotometer (Tecan Trading AG; Switzerland). One unit of enzyme activity was defined as the amount of the enzyme liberating 1 μM GlcN in 1□h at 40°C.

## 3. Results

### 3.1 CatOpt outperformed existing predictive models

First, the hyperparameter optimization on the number of attention heads (n_head) and residual dense blocks (n_RD) found that n_head=4 and n_RD=4 is the best set of hyperparameters in the search scope (**see SI, Figure S2**). Then, CatOpt, with the optimal set of hyperparameters, achieved a prediction accuracy of R2=0.479, MAE=0.607 and RMSE=0.833 on the hold-out test set (**Figure 2AB and see SI, Figure S3**). In model comparison on the same test set provided by the Zenodo repository of EpHod [13], CatOpt outperformed Seq2pHopt (R2=0.417), EpHod (R2=0.399), and OphPred (R2=0.457) (**Figure 2C**). With respect to prediction errors at *pH*_*opt*_ < 6, CatOpt had a RMSE of 1.189, close to EpHod (RMSE=1.180) and lower than OphPred (RMSE=1.393) and Seq2pHopt (RMSE=1.375) (**Figure 2D**). At *pH*_*opt*_ < 8, CatOpt had a RMSE of 1.319, slightly lower than EpHod (RMSE=1.339) and OphPred (RMSE=1.331) (**Figure 2E**). For 7 enzyme classes, CatOpt had lower RMSEs than other 3 models in oxidoreductases (EC1), hydrolases (EC3), lyases (EC4), isomerases (EC5), close RMSEs to EpHod and OphPred in transferases (EC2), ligases (EC6), and a higher RMSE than EpHod and OphPred in translocases (EC7) (**Figure 2F**). In general, CatOpt exhibited good accuracy and outperformed existing predictive models of enzyme *pH*_*opt*_.

**Figure 2.**
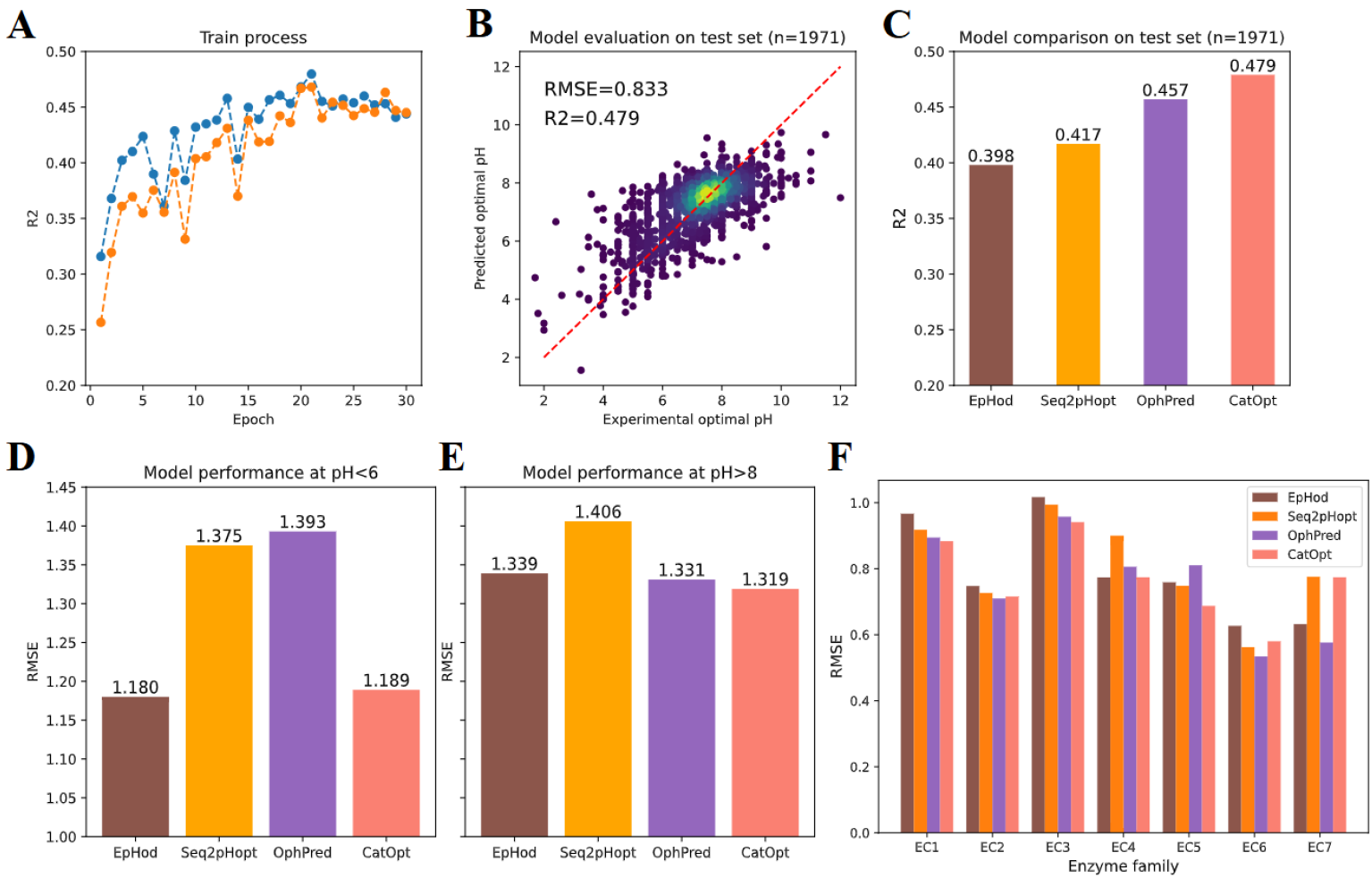
Model performance evaluation of CatOpt. (A) The R2 scores of prediction during the training process. Blue dotted curve: test set; Orange dotted curve: dev set (validation set). (B) Experimental and predicted by CatOpt (RMSE=0.833 and R2=0.479). (C) Prediction accuracy comparison of EpHod (R2=0.399), Seq2pHopt (R2=0.417), OphPred (R2=0.457), and CatOpt (R2=0.479) on the same test set. (D) Prediction accuracy comparison of 4 models at low () value range. (E) Prediction accuracy comparison of 4 models at high () value range. (F)Prediction accuracy comparison of EpHod, Seq2pHopt, OphPred, and CatOpt for different enzyme classes (EC 1-7).

### 3.2 Identifying the pH preferences of enzymes and microorganisms

After the prediction accuracy of CatOpt had been validated (**section 3.2**), this study proceeded to examine its ability to identify the pH preferences of enzymes and microorganisms. The benchmark dataset of AcalPred (54 acidic enzymes and 68 alkaline enzymes) [8] was used in the identification of acidic and alkaline enzymes. The predicted *pH*_*opt*_ values of alkaline enzymes were significantly higher than those of acidic enzymes (**Figure 3A**). With the cutoff of *pH*_*opt*_ =7.0, CatOpt classified acidic and alkaline enzymes with an accuracy of 91.8%, 50 acidic enzymes and 62 alkaline enzymes were accurately identified (**Figure 3BC**). In 10 misclassified enzymes, 4 acidic enzymes were classified as alkaline enzymes, 6 alkaline enzymes were classified as acidic enzymes. Overall, this case study demonstrated that CatOpt could discriminate enzymes with different pH preferences.

**Figure 3.**
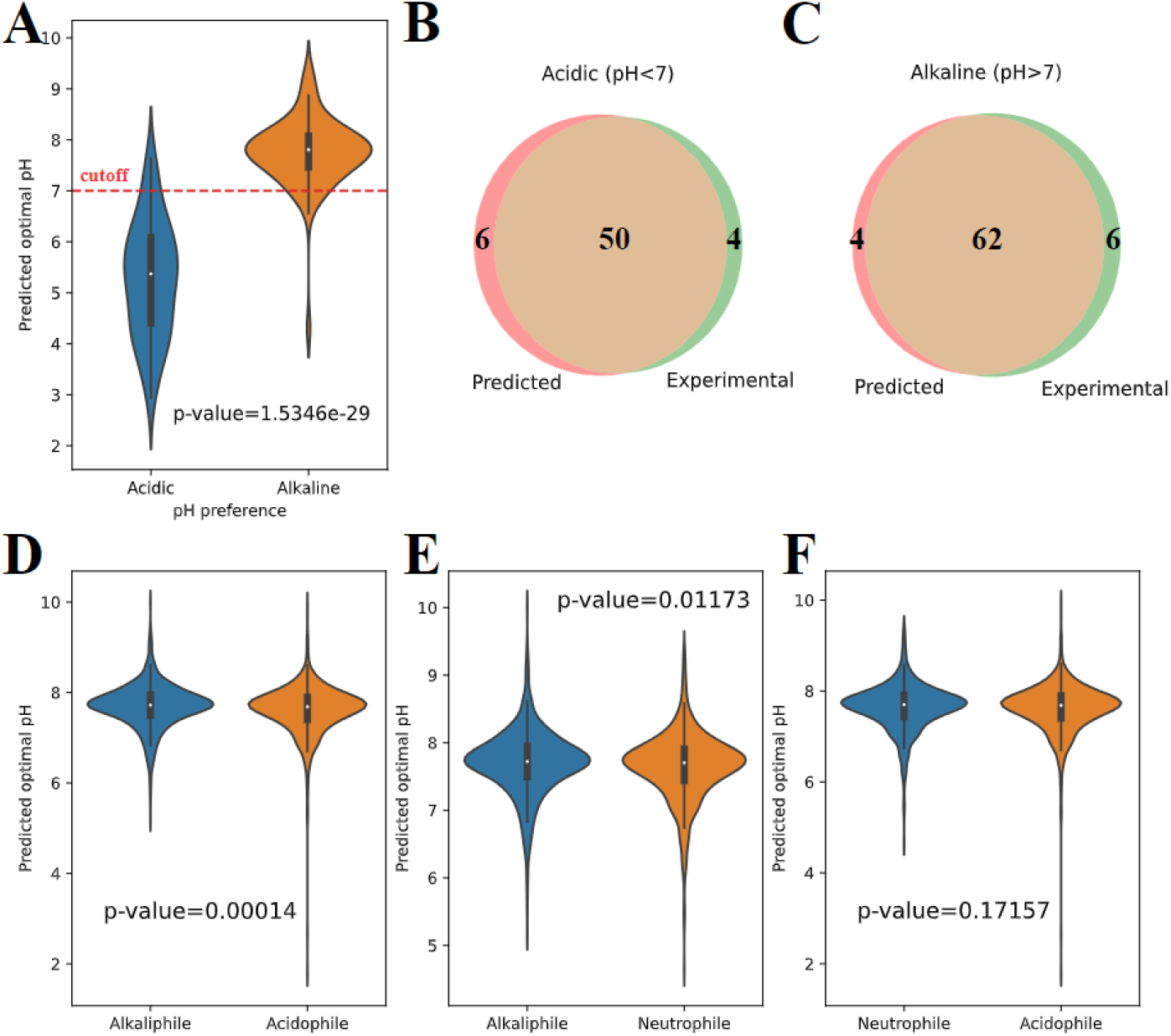
The performance of CatOpt on the pH preference of enzymes and microorganisms. (A) The distribution of predicted values of 54 acidic enzymes and 68 alkaline enzymes (p-value<0.001). (B) The venn diagram of predicted and experimental acidic enzymes. (C) The venn diagram of predicted and experimental alkaline enzymes. (D) The distribution of predicted values of enzymes from alkaliphilic and acidophilic microorganisms (p-value<0.001). (E) The distribution of predicted values of enzymes from alkaliphilic and neutrophilic microorganisms (p-value<0.05). (F) The distribution of predicted values of enzymes from neutrophilic and acidophilic microorganisms (p-value>0.05).

Next, *pH*_*opt*_ values were predicted for catalytic enzymes belonging to 3 acidophilic, 3 neutrophilic, and 3 alkaliphilic microorganisms (**see SI, Figure S4**). Enzyme protein sequences were all obtained from the uniprot database [6]. CatOpt could discriminate alkaliphilic and acidophilic microorganisms with significantly different distributions of predicted enzyme *pH*_*opt*_ values, the same for alkaliphilic and neutrophilic microorganisms (**Figure 3DE**). However, there was no significant difference between predicted *pH*_*opt*_ values of enzymes from acidophilic and neutrophilic microorganisms (**Figure 3F**). Despite that CatOpt could identify the pH preferences of enzymes, it could not accurately classify acidophilic, neutrophilic, and alkaliphilic microorganisms based on distributions of predicted enzyme *pH*_*opt*_ values.

### 3.3 Residue attention weights capture key sequence information

To investigate how residue attention weights capture important sequence information, this study compared attention weights on different types of residues across acidophilic, neutrophilic, and alkaliphilic enzymes in the hold-out test set. For both acidic and basic residues, the attention weights of acidophilic and neutrophilic enzymes were significantly higher than the weights of acidophilic enzymes (**Figure 4A**), which revealed the importance of ionizable residues on enzyme *pH*_*opt*_. In all three types of enzymes, the attention weights on non-polar residues were significantly higher than the weights on polar residues (**Figure 4B**). Next, 173 enzymes with annotated active and binding sites were selected from the test set. The active sites are regions where chemical reactions happen, and the binding sites are residues where substrates bind. The attention weights on active and binding sites were significantly higher than attention weights on other residues (**Figure 4C**), suggesting that residue attention weights could capture important residues for enzyme catalysis. For example, the active sites of 4 mannan endo-1,4-beta-mannosidases were mostly close to peaks of residue attention weights, although the weights were not indicative for binding sites (**Figure 4D**). In a nutshell, residue attention weights in CatOpt provided good interpretability by capturing key sequence information, i.e., ionizable residues, active and binding sites.

**Figure 4.**
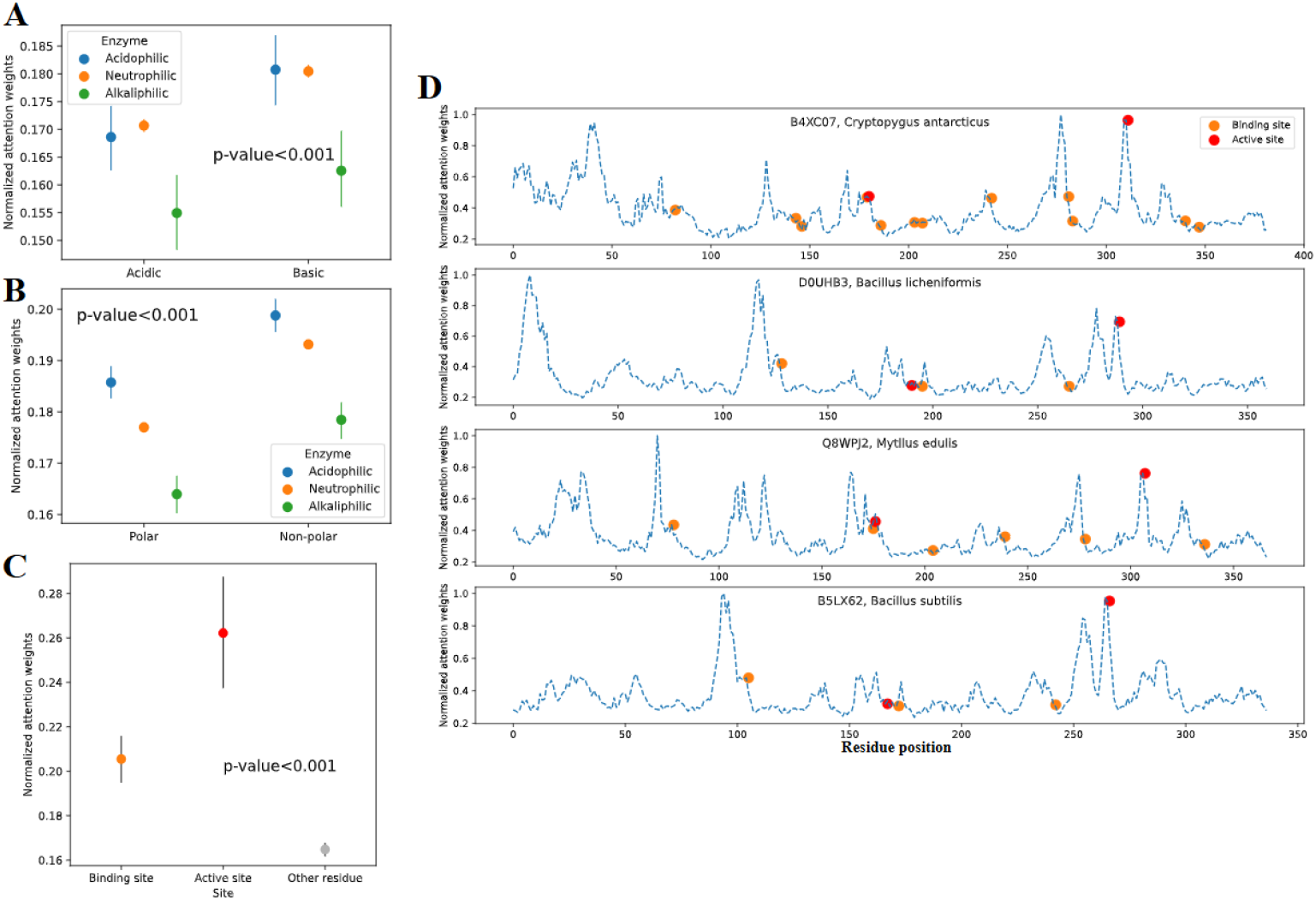
The analysis of residue attention weights. (A) The attention weights of acidic and basic residues of acidophilic, neutrophilic, and alkaliphilic enzymes. (B) The attention weights of polar and non-polar residues of acidophilic, neutrophilic, and alkaliphilic enzymes. (C) The attention weights on binding sites, active sites and other residues. (D) Representative examples of residue attention weights, positions of binding and active sites of 4 mannan endo-1,4-beta-mannosidases (EC:3.2.1.78). The uniprot IDs are B4XC07, D0UHB3, Q8WPJ2, and B5LX62. Blue dashed curve: normalized attention weights; Orange dot: binding site; Red dot: active site.

### 3.4 Prediction of optimal pH shifts caused by point mutations

To examine the inference ability of CatOpt on how point mutations affect enzyme optimal pH, this study used it to predict values of wild-types (WTs) and mutants for *Pyrococcus horikoshii* diacetylchitobiose deacetylase (PhDac) [19], *Bacillus circulans* xylanase (BCX) [25], *Lactiplantibacillus plantarum* tannase (TanBlp) [26], and *Clonostachys rosea* zearalenone hydrolase (CRZHD) [27] (**see SI, Table S1**). The experimental data of those enzymes was not included in the training set of CatOpt. The overall prediction error for WTs and mutants of those 4 different enzymes was RMSE=1.35 and MAE=1.12 (**Figure 5A**).

**Figure 5.**
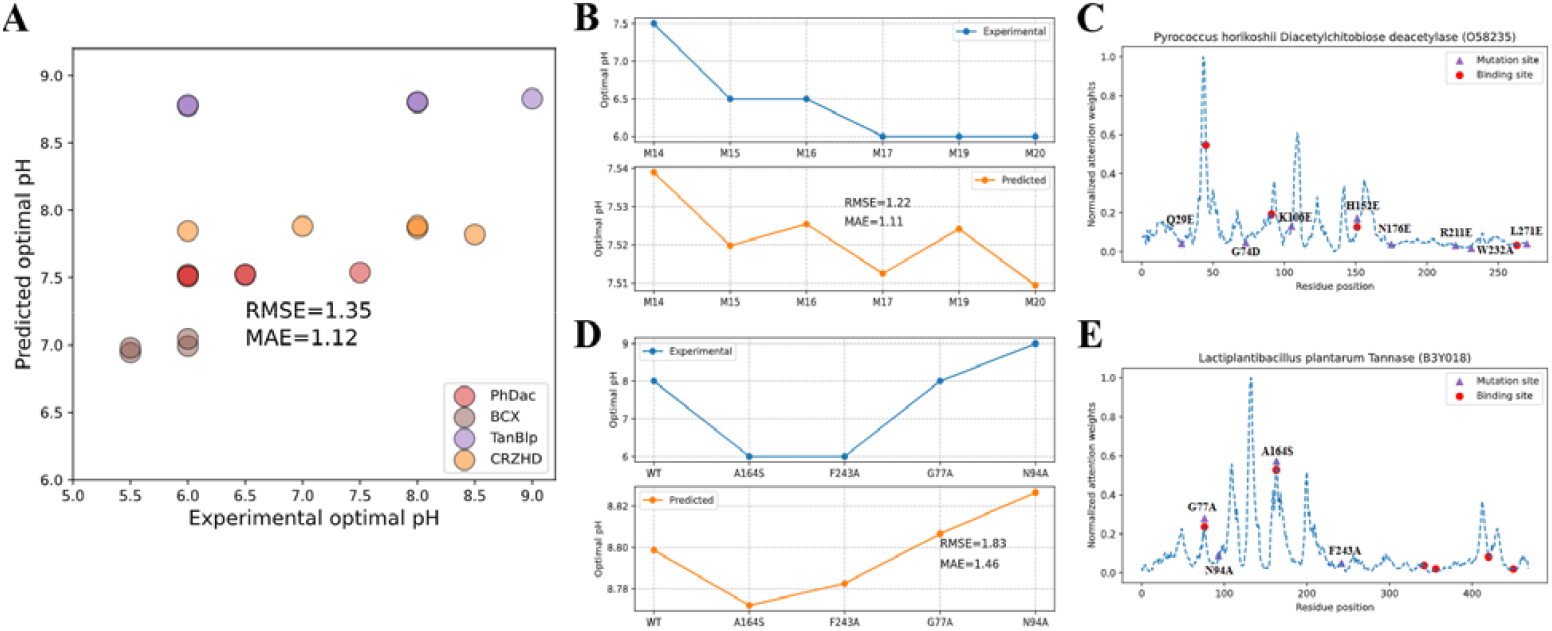
The prediction of enzyme shifts caused by mutations. (A) Experimental and predicted enzyme of WTs and mutants of 4 different enzymes (RMSE=1.35, MAE=1.12). PhDac: Pyrococcus horikoshii diacetylchitobiose deacetylase, BCX: Bacillus circulans xylanase, TanBlp: Lactiplantibacillus plantarum tannase, CRZHD: Clonostachys rosea zearalenone hydrolase, WT: wild type. (B) Experimental and predicted enzyme of 6 mutants of PhDac. (C) Residue attention weights of PhDac and positions of substrate binding sites and point mutations in 6 mutants. (D) Experimental and predicted enzyme of WT and mutants of TanBlp. (E) Residue attention weights of TanBlp and positions of substrate binding sites and point mutations in 4 mutants.

For PhDac, the prediction by CatOpt qualitatively accounted for the downshift of *pH*_*opt*_ in M15 and M16 in comparison to M14, and in M17 and M20 in comparison to M15 and M16, but the numerical difference of predicted *pH*_*opt*_ among 6 mutants was small (**Figure 5B**). The residue attention weights of PhDac captured substrate binding sites with high peaks (e.g., D46) (**Figure 5C**), although the spatial distribution of weights could not explain effective point mutations causing down-shift of *pH*_*opt*_ (e.g., Q29E). The highest peak of attention weights was at residue 44, suggesting a potential effective site for point mutations. For TanBlp, CatOpt quantitatively predicted the downshift of *pH*_*opt*_ caused by A164S and F243A in comparison to WT, and the upshift of *pH*_*opt*_ by G77A and N94A in comparison to A164S and F243A (**Figure 5D**). Two substrate binding sites of TanBlp (G77 and A164) and two effective point mutations (G77A and A164S) were identified by high peaks of residue attention weights (**Figure 5E**). BCX and CRZHD were not included in further analysis (**see SI, Figure S6**), due to lack of protein sequence annotation. Generally speaking, this case study demonstrated CatOpt’s capability to predict the effect of point mutations on enzyme *pH*_*opt*_, although the numerical differences of predicted enzyme *pH*_*opt*_ for WTs and mutants were relatively smaller than those observed in experimental measurements.

### 3.5 Predictor-guided engineering of diacetylchitobiose deacetylase to enhance acid resistance

CatOpt was used as a computational design tool to enhance the acid resistance of *Pyrococcus horikoshii* diacetylchitobiose deacetylase (PhDac) (**Figure 5BC**), which is used in the environmentally-friendly manufacturing of GlcN [19]. Enzyme and turnover number values of 1197 mutants with single point mutations based on M20 (**see SI, Table S1**) were predicted (**Figure 6A**). 5 mutants with lowest predicted values, which also had relatively high turnover number values, were selected to examine their activities at pH=4.5 and 5.5. Compared to M20, H44C improved the activities of PhDac at pH=4.5 and 5.5 by around 7%, H44D improved the activity at pH=4.5 by around 2% (**Figure 6B**). The other 3 selected mutants (M20+H44I, M20+H44N, M20+H44P) did not enhance the acid resistance, but their activities at pH=4.5 were all higher than activities at pH=5.5 (**Figure 6B**), which suggested the downshift of. Both effective point mutations (H44C and H44D) substituted a basic residue (Histidine (H)) with an acidic residue (Cysteine (C) and Aspartate (D)). In addition, enzyme activities of 5 un-selected mutants (**Figure 6A**) were also measured, and they showed weaker acid resistance than M20 (**Table S2, 3, Figure S7**). In short, the success of computationally designed point mutations showed the usefulness of CatOpt as a design tool of enzyme engineering.

**Figure 6.**
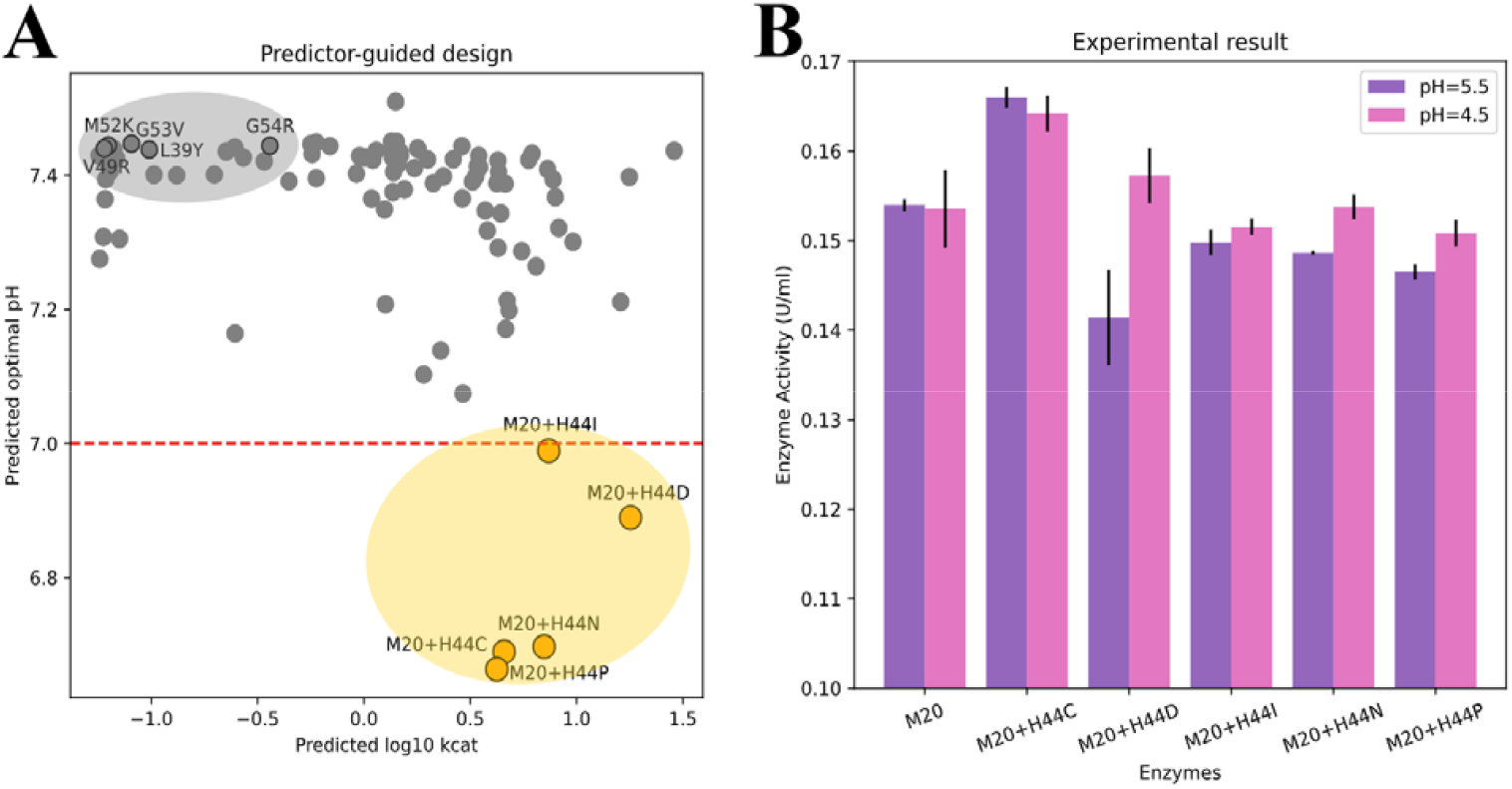
Predictor-guided engineering of Pyrococcus horikoshii diacetylchitobiose deacetylase (PhDac). (A) Predicted enzyme and turnover number values () of mutants with single point mutations based on M20 (Only lower 10% of predicted values were presented). Orange dots represent 5 selected mutants: M20+H44I, M20+H44D, M20+H44N, M20+H44C, M20+H44P. Grey dots represent the remaining un-selected mutants, grey dots with black edges represent M20+G53V, M20+M52K, M20+G54R, M20+V49R, M20+L39Y. (B) Enzyme activities (U/ml) at pH=4.5 and 5.5 for M20 and 5 selected mutants. All measurements have 3 replicates (**see SI, Table S2, 3**). The error bars represent standard deviations.

## 4. Discussion

To address the limitations of existing predictive models of *pH*_*opt*_ (e.g., low accuracy or lack of interpretability), this study developed CatOpt, a deep learning model capable of predicting enzyme *pH*_*opt*_ directly from protein sequences. Main components of CatOpt were ESM-2 embedding generation, multi-scale CNN, multi-head self-attention, and residual dense neural networks (**Figure 1**). Compared with one-hot encoding [28] or k-mer based dictionary embedding [23,29], ESM-2, as a pre-trained large language model of proteins, can transfer the knowledge of protein structures and functions from a large dataset of millions of protein sequences to this prediction task using thousands of protein sequences [16]. The advantage of multi-scale CNN with different window sizes over CNN with a fixed window size lies in ensembling information to enrich the representation of protein sequence features [30]. In contrast to multi-head light attention in Seq2Topt/Seq2pHopt [14], self-attention in CatOpt can model dependencies between different regions of the protein sequence [31]. Additionally, the use of residue dense neural networks instead of multiple linear layers could effectively reduce the vanishing and exploding gradient issues in deep neural networks [32]. Consequently, CatOpt outperformed existing enzyme *pH*_*opt*_ predictors (e.g., OphPred) with RMSE =0.833 and R2=0.479, and provided good interpretability with informative residue attention weights (**section 3.3**).

The classification of acidic/alkaline enzymes (**section 3.2**), prediction of enzyme *pH*_*opt*_ shifts by point mutations (**section 3.4**), and predictor-guided engineering of PhDac (**section 3.5**) demonstrated that CatOpt could be applied to enzyme mining and computational design of enzymes via fast screening the effect of point mutations. Besides in-silico screening, the combination of generative deep learning and CatOpt might lead to automatic generation of novel acidic or alkaline enzymes through predictor-guided generator optimization [33]. Moreover, the informative attention weighted protein features extracted by CatOpt (**section 3.3**) could potentially be used to improve the performances of other prediction tasks, such as the prediction of pH-dependent enzyme turnover numbers [34,35], via transfer learning.

Despite the achievement of CatOpt outlined above, some limitations still exist and hinder its performance. Similar to Seq2Topt/Seq2pHopt [14], the accuracy of CatOpt was also affected by the imbalance of the training dataset. Oversampling and loss reweighting can mitigate the imbalance, but the prediction accuracy at low and high value ranges will still be relatively low [13,14]. Using high-throughput enzyme assays to append entries at the ranges of *pH*_*opt*_ <6 and *pH*_*opt*_ >8 to the dataset is necessary to further improve the performance of *pH*_*opt*_ prediction. The neglect of environmental factors influencing enzyme *pH*_*opt*_ is another shortcoming, such as temperature [36]. Although the inclusion of metadata might improve the prediction accuracy, a large portion of enzyme assay results curated from databases miss such information. In the attempt to identify organismal pH preferences (**section 3.2**), CatOpt failed to differentiate neutrophilic and acidophilic microorganisms with distributions of predicted enzyme *pH*_*opt*_ values. The distribution of experimental enzyme *pH*_*opt*_ values indicates that most enzymes in acidophilic microorganisms have *pH*_*opt*_ in the range of 6∼8, just like neutrophilic microorganisms (**see SI, Figure S5**). Therefore, the relationship between microbial growth *pH*_*opt*_ and enzyme *pH*_*opt*_ still remains to be investigated.

In conclusion, CatOpt is an interpretable deep learning predictor of enzyme *pH*_*opt*_ that demonstrated improved accuracy compared to existing tools, despite the limitations discussed above. As envisaged, CatOpt can potentially accelerate enzyme discovery for desired properties from “biological dark matter” and enzyme engineering with *in-silico* design.

## Acknowledgements

This work was financially supported by the National Key Research and Development Program of China (No. 2023YFA0914500), and the National Natural Science Foundation of China (No. 32171261). The authors would like to acknowledge the use of the University of Oxford Advanced Research Computing (ARC) facility (http://dx.doi.org/10.5281/zenodo.22558) in carrying out this work.

## Author contributions

Sizhe Qiu constructed the deep learning model, designed point mutations, and produced the first draft. Nankai Wang conducted the experiment and contributed to the first draft. Yishun Lu assisted in model construction and contributed to model optimization. Jinsong Gong participated in the writing and review of the first draft. Aidong Yang and Jinsong Shi supervised this research project and critically reviewed the manuscript.

## Abbreviations

CNN: convolutional neural network
GlcN: glucosamine
GlcNAc: N-acetylglucosamine
Leaky ReLU: leaky rectified linear unit
MAE: mean absolute error
MSE: mean squared error
PhDac: *Pyrococcus horikoshii* diacetylchitobiose deacetylase
*pH*_*opt*_: optimal pH
RD: residual dense block
R2, r-squared: the coefficient of determination
RMSE: root mean squared error
WT: wild-type.

## Data availability statement

The code and data are openly available at https://github.com/SizheQiu/CatOpt and supplementary information.

