## Supplementary material for "Deep learning-based prediction of enzyme optimal pH and design of point mutations to improve acid resistance": Figure S1-7, Table S1-3

### Authorship

Sizhe Qiu^1‡^, Nan-Kai Wang^2,3‡^, Yishun Lu^1,4‡^, Jin-Song Gong^2,3*^, Jin-Song Shi^2,3^, Aidong Yang^1^*

^1^Department of Engineering Science, University of Oxford, OX1 3PJ, United Kingdom

^2^Key Laboratory of Carbohydrate Chemistry and Biotechnology, Ministry of Education, School of Life Sciences and Health Engineering, Jiangnan University, Wuxi, 214122, PR China

^3^Yixing Institute of Food and Biotechnology Co., Ltd, Yixing, 214200, PR China

^4^Oxford e-Research Centre, Department of Engineering Science, University of

Oxford, 7 Keble Road, Oxford, United Kingdom

^‡^Equal contribution

### 1. Supplementary Methods

#### 1.1 Software and code availability

All scripts were written in python. The deep learning model was implemented using PyTorch v1.12.0. The computer used in this work was a Dell Latitude Laptop with intel core i7 CPU. The model was trained with GPU RTX8000 provided by Advanced Research Computing (ARC) service in the University of Oxford [[1]](https://paperpile.com/c/LH39uq/pSS1). Figures in the main text and supplementary information were all edited using InkScape (<https://inkscape.org/>).

#### 1.2 Evaluation metrics

To quantitatively assess the prediction accuracy, R2 (Eq. S1), RMSE (Eq. S2), and MAE (Eq. S3) were computed for each test.

$R^{2}=\frac{\sum_{i=1}^{n} (y_{ie}-y_{ip})^{2}}{\sum_{i=1}^{n} (y_{ie}-\underline{y})^{2}} (Eq. S1)$

$RMSE=\sqrt{\frac{1}{n}\sum_{i=1}^{n} (y_{ie}-y_{ip})^{2}} (Eq. S2)$

$MAE = \frac{1}{n}\sum_{i=1}^{n} \left| y_{ie}-y_{ip} \right| (Eq. S3)$

### 2. Supplementary Figures

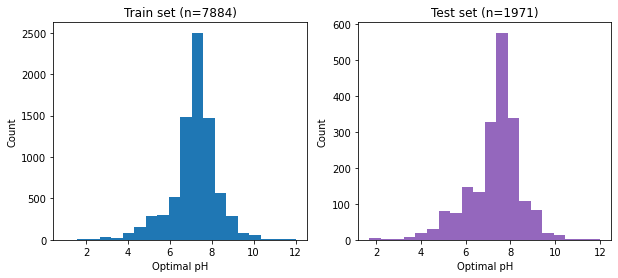

Figure S1. The distributions of enzyme optimal pH values in the train set (7884 entries) and test set (1971 entries).

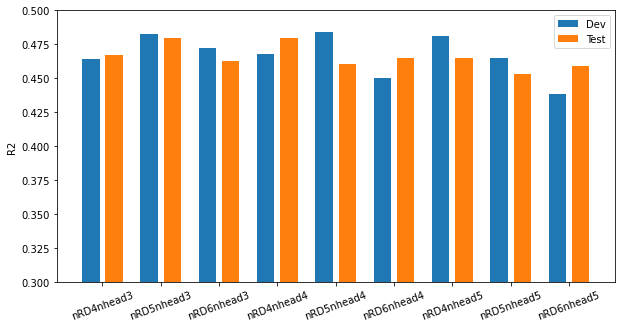

Figure S2. Hyperparameter optimization of enzyme optimal pH prediction for number of attention heads (n_head) and residual dense blocks (n_RD). n_RD=4, n_head=4 was chosen as the best set of hyperparameters.

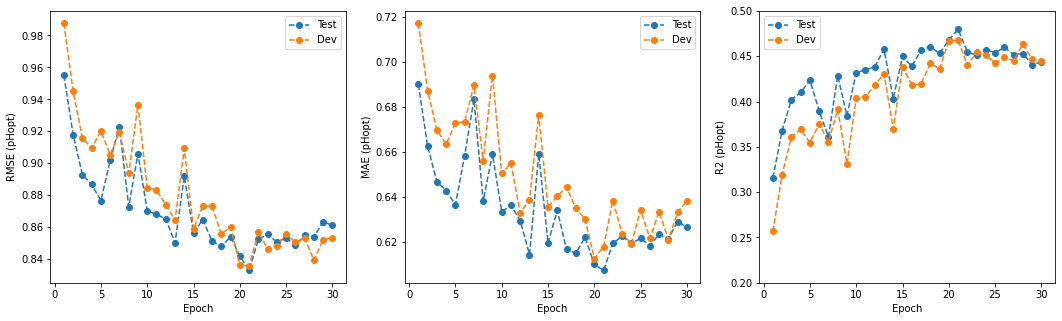

Figure S3. The RMSE, MAE, R2 scores of CatOpt during the training process.

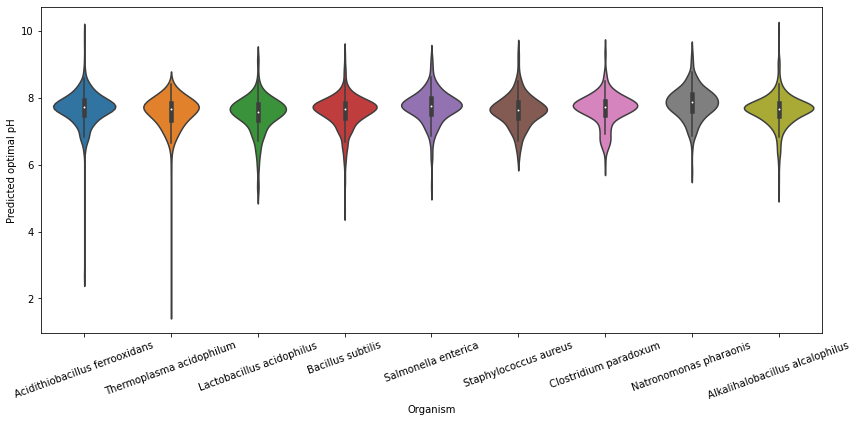

Figure S4. Predicted optimal pH values for enzymes in 9 microorganisms. Acidophiles: *Acidithiobacillus ferrooxidans*, *Lactobacillus acidophilus*, and *Thermoplasma acidophilum*; Neutrophiles: *Bacillus subtilis*, *Salmonella enterica*, and *Staphylococcus aureus*; Alkaliphiles: *Clostridium paradoxum*, *Alkalihalobacillus alcalophilus*, and *Natronomonas pharaonis*.

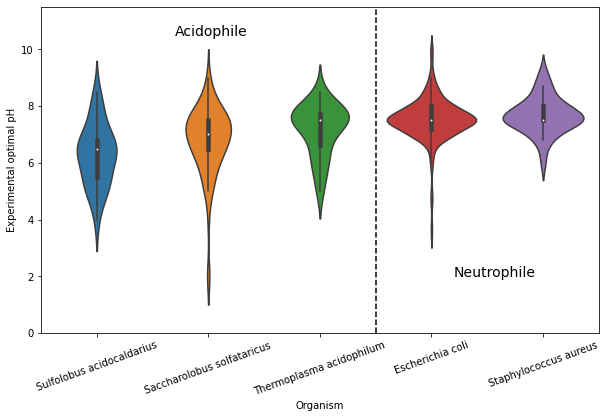

Figure S5. Experimental optimal pH values for enzymes in 3 acidophilic microorganisms and 2 neutrophilic microorganisms.

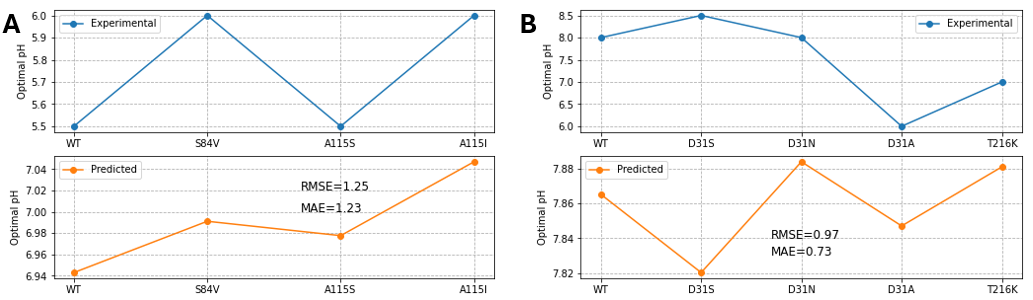

Figure S6. (A) Experimental and predicted enzyme $pH_{opt}$ of wild-types and mutants of BCX. (B) Experimental and predicted enzyme $pH_{opt}$ of wild-types and mutants of CRZHD.

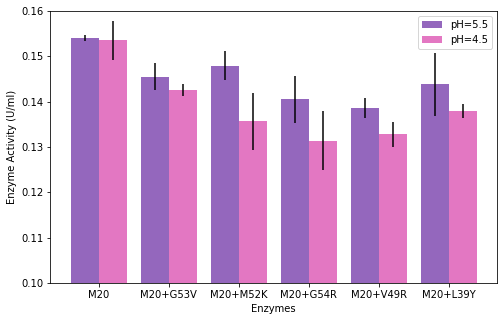

Figure S7. Enzyme activities (U/ml) at pH=4.5 and 5.5 for M20 and M20+G53V, M20+M52K, M20+G54R, M20+V49R, M20+L39Y.

### 3. Supplementary Tables

Table S1.Information of wild-type and mutated enzymes used in case studies.

| Enzyme | Mutations | $pH_{opt}$ | Reference |
| --- | --- | --- | --- |
| *Pyrococcus horikoshii* diacetylchitobiose deacetylase | M14: G74D/H152E/W232A | 7.5 | [[2]](https://paperpile.com/c/LH39uq/XHBi) |
|  | M15: G74D/H152E/W232A/Q29E | 6.5 |  |
|  | M16: G74D/H152E/W232A/Q29E/K106E | 6.5 |  |
|  | M17: G74D/H152E/W232A/Q29E/K106E/N176E | 6.0 |  |
|  | M19: G74D/H152E/W232A/Q29E/K106E/N176E/R221E | 6.0 |  |
|  | M20: G74D/H152E/W232A/Q29E/K106E/N176E/R221E/L271E | 6.0 |  |
| *Bacillus circulans* xylanase | WT | 5.5 | [[3]](https://paperpile.com/c/LH39uq/ZqJg) |
|  | S84V | 6 |  |
|  | A115S | 5.5 |  |
|  | A115I | 6 |  |
| *Clonostachys rosea* zearalenone hydrolase | WT | 8 | [[4]](https://paperpile.com/c/LH39uq/t3ga) |
|  | D31S | 8.5 |  |
|  | D31N | 8 |  |
|  | D31A | 6 |  |
|  | T216K | 7 |  |
| *Lactiplantibacillus plantarum* tannase | WT | 8 | [[5]](https://paperpile.com/c/LH39uq/pE5w) |
|  | A164S | 6 |  |
|  | F243A | 6 |  |
|  | G77A | 8 |  |
|  | N94A | 9 |  |

Table S2. The measured enzyme activities (U/ml) of PhDac mutants at pH=5.5.

| Mutant ID | Replicate 1 | Replicate 2 | Replicate 3 |
| --- | --- | --- | --- |
| M20 | 0.1543 | 0.1532 | 0.1544 |
| M20+H44C | 0.1649 | 0.1658 | 0.1672 |
| M20+H44D | 0.1454 | 0.1435 | 0.1354 |
| M20+H44I | 0.1508 | 0.1504 | 0.1482 |
| M20+H44N | 0.1484 | 0.1488 | 0.1487 |
| M20+H44P | 0.1456 | 0.1468 | 0.1472 |
| M20+G53V | 0.1455 | 0.1484 | 0.1426 |
| M20+M52K | 0.1492 | 0.1443 | 0.1503 |
| M20+G54R | 0.1369 | 0.1381 | 0.1465 |
| M20+V49R | 0.1404 | 0.1390 | 0.1362 |
| M20+L39Y | 0.1514 | 0.1423 | 0.1378 |

Table S3. The measured enzyme activities (U/ml) of PhDac mutants at pH=4.5.

| Mutant ID | Replicate 1 | Replicate 2 | Replicate 3 |
| --- | --- | --- | --- |
| M20 | 0.1517 | 0.1505 | 0.1585 |
| M20+H44C | 0.1629 | 0.1665 | 0.1632 |
| M20+H44D | 0.1552 | 0.1558 | 0.1608 |
| M20+H44I | 0.1516 | 0.1506 | 0.1524 |
| M20+H44N | 0.1534 | 0.1526 | 0.1553 |
| M20+H44P | 0.1494 | 0.1523 | 0.1508 |
| M20+G53V | 0.1426 | 0.1440 | 0.1412 |
| M20+M52K | 0.1328 | 0.1313 | 0.1429 |
| M20+G54R | 0.1355 | 0.1239 | 0.1347 |
| M20+V49R | 0.1306 | 0.1319 | 0.1358 |
| M20+L39Y | 0.1393 | 0.1381 | 0.1364 |
